# Expression of elastin, FBXW2, fibrillin-1, and α-smooth muscle actin in damaged blood vessels of bovine legs

**DOI:** 10.1101/2024.02.06.579234

**Authors:** Mari Akiyama

## Abstract

Elastic fibers present within the internal and external elastic laminae play a vital role in maintaining the health of blood vessels. The impairment of smooth muscle cell function results in the development of vascular diseases. In this study, blood vessels from bovine legs were carefully examined in culture dishes for 5 weeks. Examination of elastin, F-box and WD-40 domain-containing protein 2 (FBXW2), and fibrillin-1 in serial sections revealed that elastin and FBXW2 were present in the internal and external elastic laminae, whereas fibrillin-1 was present outside of them. After 5 weeks, elastin and FBXW2 maintained the shape of the elastic lamina, whereas the fibrillin-1-rich layer spread and migrated from the elastic lamina. When the blood vessel lumen was blocked by tissue rich in fibrillin-1, α-smooth muscle actin (SMA) expression weakened. Although elastin and fibrillin-1 are components of elastic fibers, this study revealed that fibrillin-1 has a role different from that of elastin. This in vitro model is useful for identifying the mechanisms underlying vascular degradation.

## Introduction

Vascular diseases, such as atherosclerosis [1] and aortic aneurysm [2], are life-threatening, and the aging process of elastic fibers affect cardiovascular disease development [3] and inhibition of smooth muscle cell proliferation increases inflammatory effects [4]. Isolated smooth muscle cells have been examined in a study on cardiovascular diseases [5]. Fibrillin microfibrils are elastic fiber components [3] [6] . Mutated fibrillin-1 causes Marfan syndrome [7] [8], and the relationship between fibrillin-1-derived asprosin and metabolic disorders remains controversial [9].

In 2022, Akiyama [10] reported that both elastin and F-box and WD-40 domain-containing protein 2 (FBXW2) were expressed in the elastic fibers of the bovine periosteum and blood vessels. However, the relationship between elastic fibers and FBXW2 in blood vessels remains unclear. In this study, the extent of damage to the elastic lumen, elastic fibers, and vessel walls over a 5-week culture period was investigated. The aim of this study was to clarify whether changes in elastic fibers and smooth muscle cells are involved in blood vessel damage.

## Materials and Methods

### Serial sections

The use of blood vessels was approved by the Osaka Dental University Regulations on Animal Care and Use (Approval No. 23-02007). Importantly, no live cows were used in this study. In addition, this research did not involve human participants.Four blood vessels from randomly selected legs of 30-month-old Japanese Black Cattle (Kobe Chuo Chikusan, Kobe, Japan) were dissected into three parts.

Blood vessels from the same cows were divided into three groups:

a. day 0;
b. five-week culture in Medium 199 (12340-030; Gibco, Grand Island, NY, USA) with 5 mg/mL ascorbic acid (A0278-25G; Sigma, St. Louis, MO, USA); and
c. five-week culture in Medium 199 (12340-030; Gibco) without ascorbic acid.

Medium 199 contained fetal bovine serum and antibiotics, as previously described [10]. On day 0 or after 5 weeks, blood vessels were fixed in 4% paraformaldehyde, and paraffin-embedded sections were obtained.

### Immunohistochemistry

For antigen retrieval, Proteinase K (S3020; Dako Cytomation, Glostrup, Denmark) was used for the antibodies of elastin, FBXW2, and fibrillin-1. Tris/EDTA buffer (pH 9.0, 40 min; Diagnostic BioSystems, Pleasanton, CA, USA) was used for the antibody of α-smooth muscle actin (SMA). Antibodies of elastin and FBXW2 were the same as those previously described (10), but different cows were used in this study. To investigate components of elastic fibers, antibodies of elastin (dilution 1:300), FBXW2 (1:100), and fibrillin-1 (1:100; 11C1.3; GeneTex, Inc., CA, USA) were used. As a marker of vascular smooth muscle cells, the antibody of α-SMA (1:500; ARG66381; Arigo Biolaboratories Corp., Hsinchu City, Taiwan) was used. As secondary antibodies, alkaline phosphatase-conjugated AffiniPure goat anti-rabbit IgG(H+L) (SA00002-2; Proteintech Group, Inc., IL, USA), alkaline phosphatase-conjugated AffiniPure goat anti-mouse IgG(H+L) (SA00002-1; Proteintech Group), and anti-goat IgG-AP (sc-2355; Santa Cruz Biotechnology, Inc., CA, USA) were used. All images were visualized with Perma Blue/AP (K058; Diagnostic BioSystems, Pleasanton, CA, USA) under a fluorescent microscope (BZ-X810; Keyence Japan, Osaka, Japan). X800 Viewer (Keyence) and X800 Analyzer software (Keyence) were used for all images. Quantitative and statistical analyses were not performed.

## Results

Fig 1A and B presents a comparison of elastin, FBXW2, and fibrillin-1 through serial section examination. Elastin and FBXW2 were expressed in both internal and external elastic laminae, whereas fibrillin-1 exhibited weak expression within these laminae, but showed high expression outside them. After a 5-week culture, with and without ascorbic acid, FBXW2 associated with elastin and fibrillin-1 expression became prominent in the thick adventitia (Fig 2A and B). Notably, fibrillin-1 was clearly expressed in the adventitia and regions of elastic fiber cleavage. Serial sections revealed that a 5-week culture led to the blockage of blood vessel lumen (Fig 3). On day 0, no blockage was observed (Fig 3A). In a 5-week culture with ascorbic acid, elastic fiber cleavage and lumen blockage were evident (Fig 3B). In the intima layer, the expression of α-SMA weakened, and the tissue blocking the lumen contained fibrillin-1 (Fig 3B). A 5-week culture without ascorbic acid revealed that, in the region of elastic fiber cleavage, fibrillin-1-rich tissue passed from the adventitia, through the media, to the intima layer. In this region, the expression of α-SMA was weak (Fig 3C). Additionally, a comparison between fibrillin-1 and α-SMA showed that weak α-SMA expression correlated with fibrillin-1 expression in the blood vessel lumen (Fig 4).

**Fig 1.**
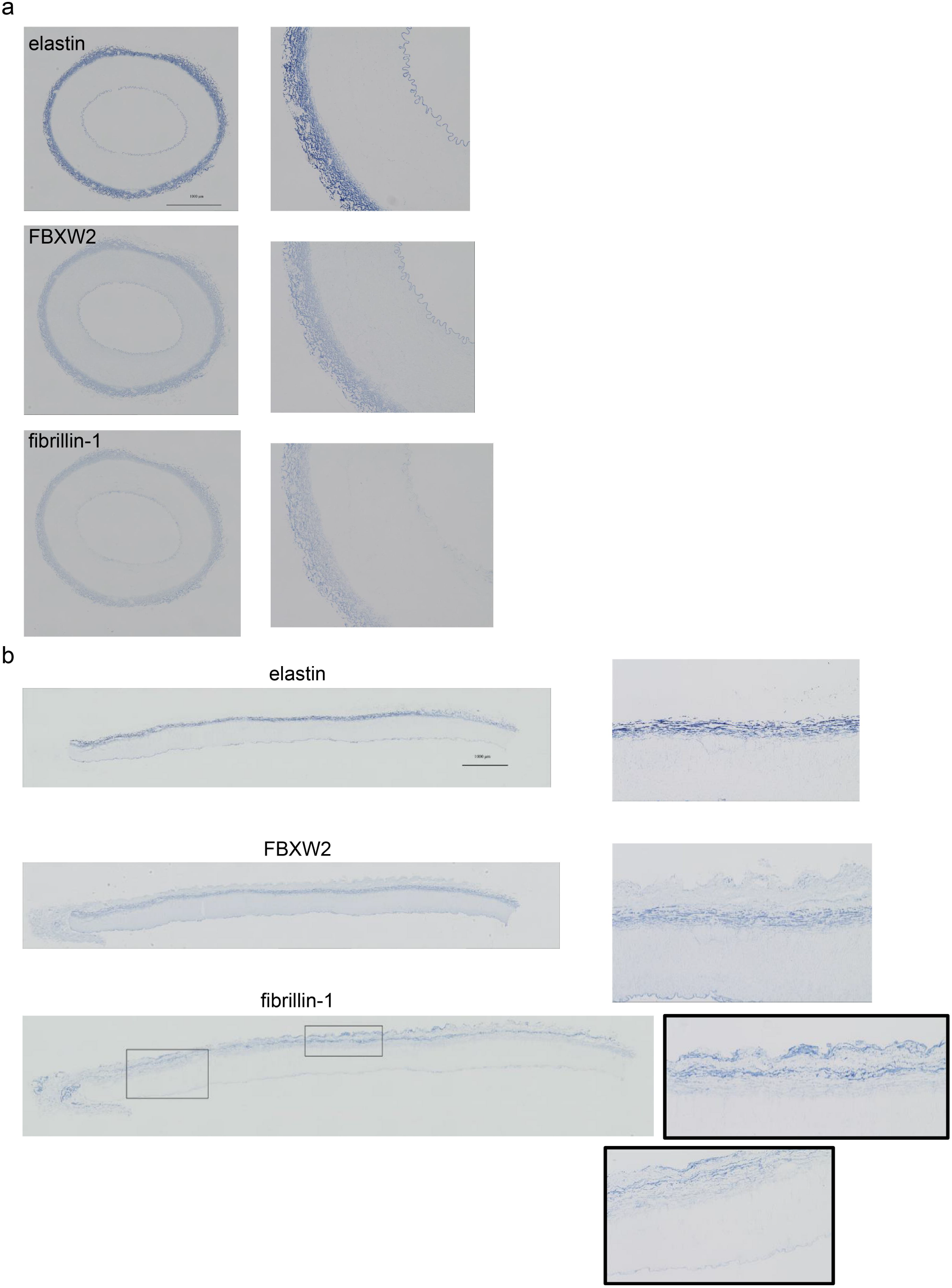
Serial sections: (top) elastin, (middle) FBXW2, (bottom) fibrillin-1 in day 0 blood vessel. (A and B) (right) High-magnification images of images on the left. Elastin and FBXW2 expression was apparent in the internal and external elastic laminae. Fibrillin-1 expression was weak in the internal and external elastic laminae, but apparent outside the external elastic lamina. Scale bar = 1000 μm.

**Fig 2.**
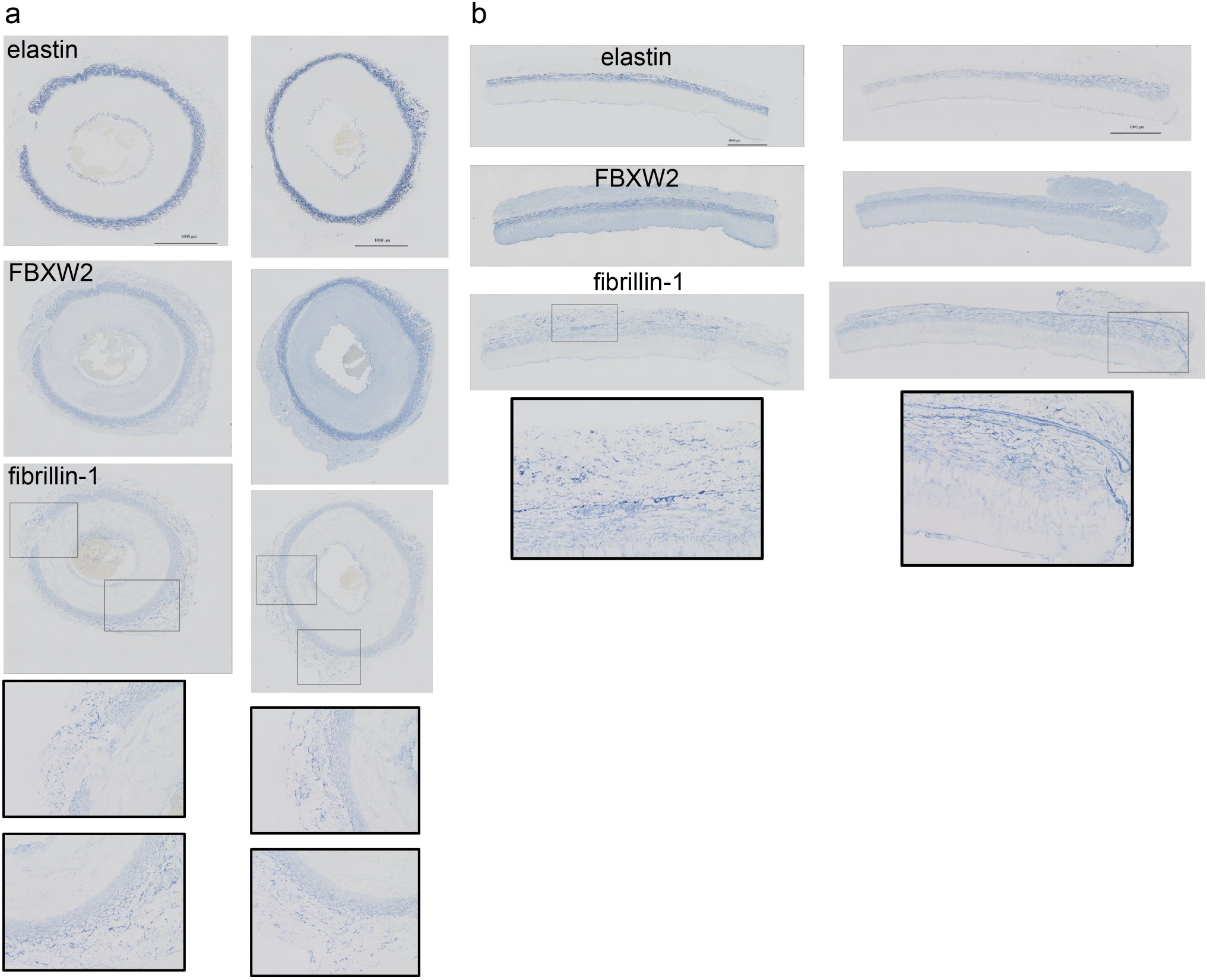
Serial sections: (top) elastin, (middle) FBXW2, (bottom) fibrillin-1 in week 5 blood vessel. (A and B) (left) With ascorbic acid; (right) without ascorbic acid. Fibrillin-1-rich adventitia becomes thick. Black-lined squares are high-magnification fibrillin-1 images. Scale bar = 1000 μm.

**Fig 3.**
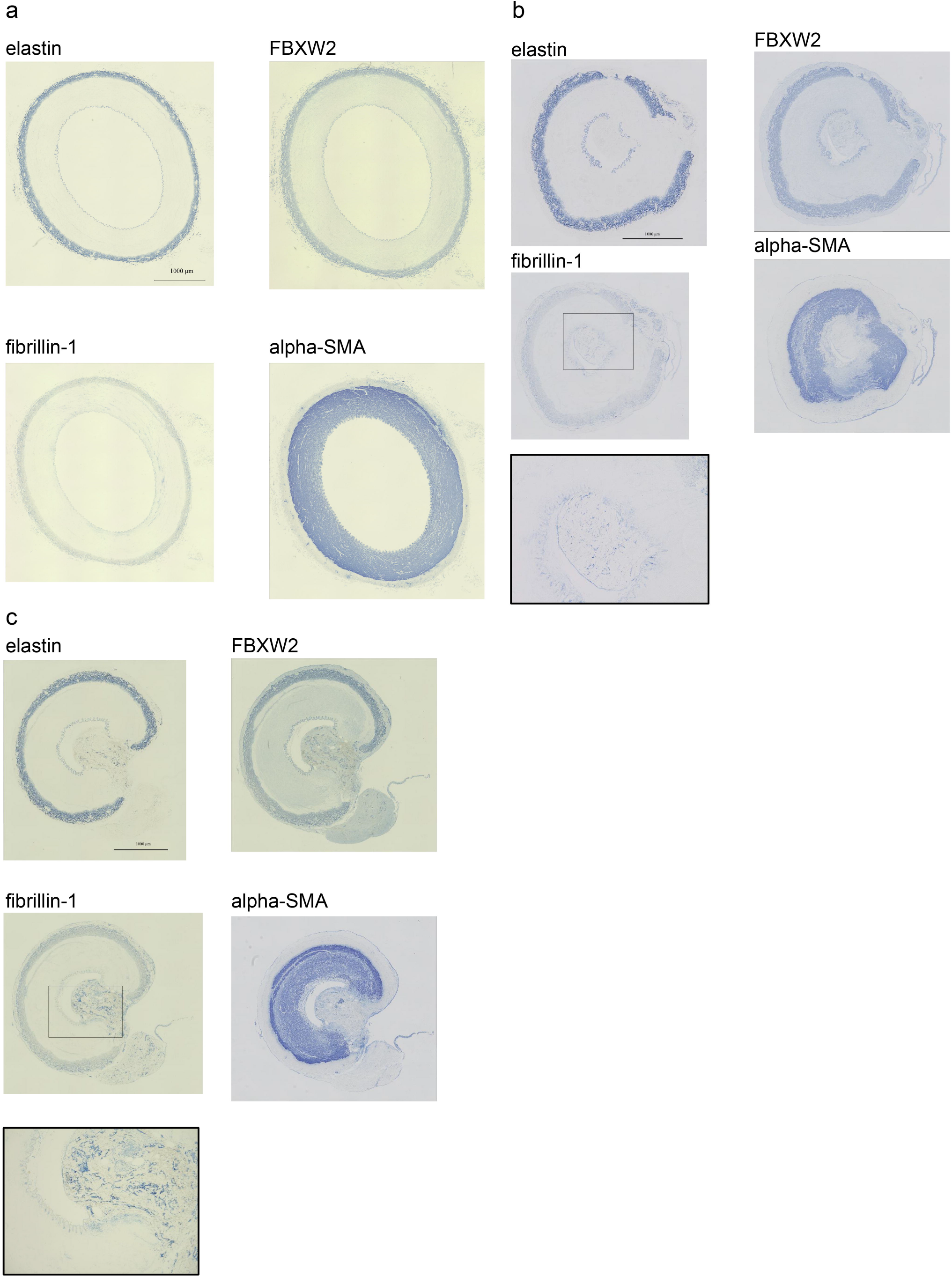
Serial sections were examined for elastin, FBXW2, fibrillin-1, and α-SMA expression. (A) Day 0. (B) Five-week culture with ascorbic acid. (C) Five-week culture without ascorbic acid. Fibrillin-1-rich tissue blocked blood vessel lumen, and α-SMA expression was weak in this region. Black-lined square is a high-magnification fibrillin-1 image. Scale bar = 1000 μm.

**Fig 4.**
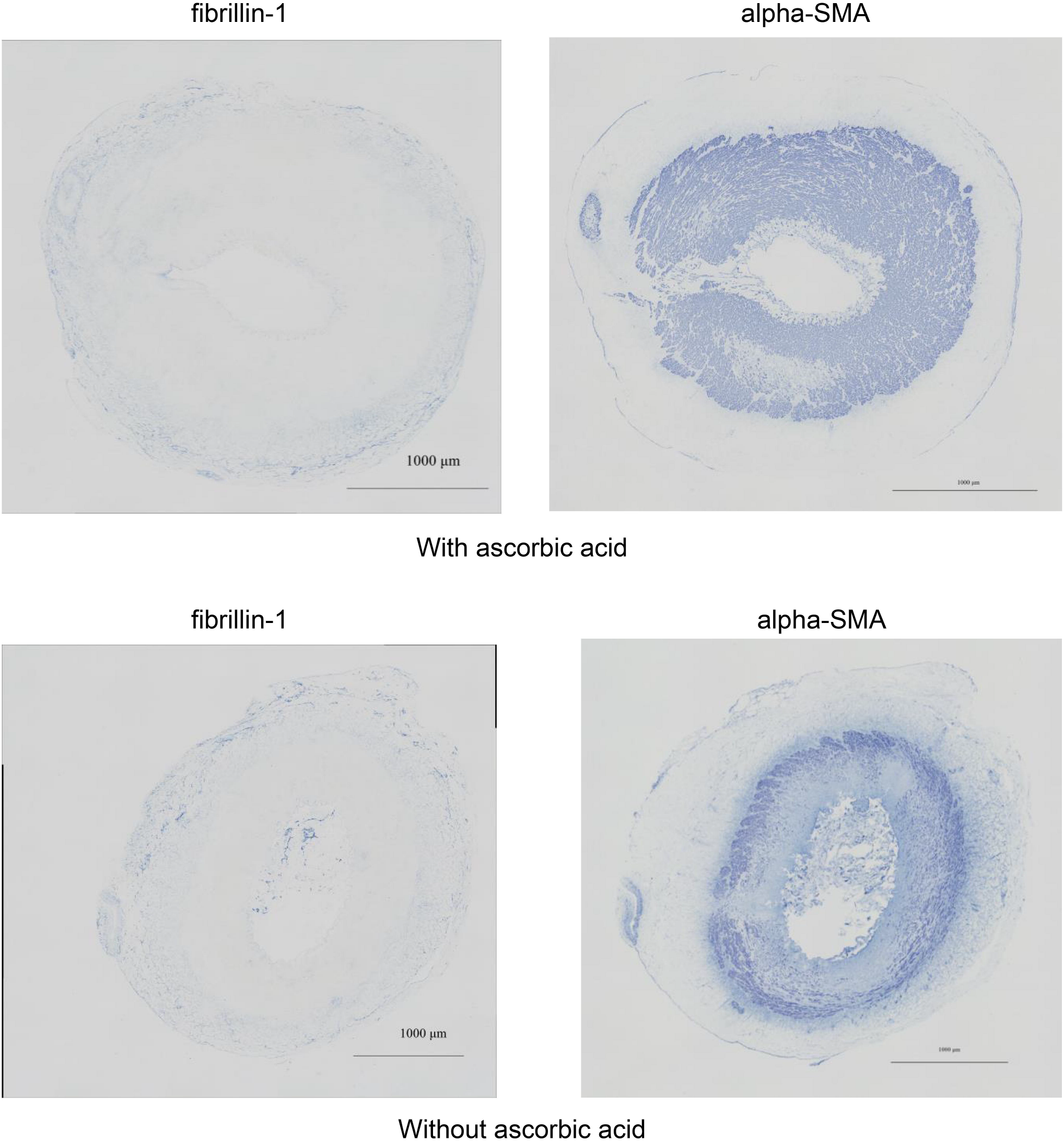
Five-week culture; comparison of fibrillin-1 and α-SMA expression. The sections are not serial sections but are from the same cows. Without ascorbic acid, fibrillin-1-rich tissue blocked the lumen and α-SMA expression became weak. Scale bar = 1000 μm.

## Discussion

Whether elastic fiber-associated FBXW2 is a component of elastic fibers remains controversial [10]. Akiyama reported the isolation of FBXW2 from elastic fibers of bovine periosteum after a 5-week culture without ascorbic acid [11]. In the present study, fibrillin-1 was expressed in a region different from that of FBXW2 (Fig 1B). These results indicate that the FBXW2 antibody did not cross-react with either elastin or fibrillin-1. This study demonstrated that from day 0 to week 5, FBXW2 did not disassociate from elastin in the elastic fibers of the blood vessels.

Although both elastin and fibrillin-1 are recognized components of elastic fibers [3, 12], fibrillin-1 expression is weak in the elastic lamina but pronounced in the adventitia (Fig 1A and B). This can be attributed to the covering by elastin or a small amount of fibrillin-1 in the elastic lamina. Steijns et al. [13] reported that, in the vessel wall, fibrillin-1 and elastic fiber were present in the tunica media, but they did not use an antibody for elastin; furthermore, small vessels were used. The present study showed that elastin and fibrillin-1 were not expressed in exactly the same region. After 5 weeks, the adventitia and fibrillin-1-rich layer thickened. It is possible that the original layer, strongly positive for fibrillin-1 on day 0 (Fig 1B), grew along with the adventitia after 5 weeks. A comparison of elastin and fibrillin-1 showed that elastin maintained the shape of lamina after 5 weeks even after cleavage, whereas the fibrillin-1-rich layer spread and migrated from the elastic lamina (Fig 2A and B).

In 2011, Bastos et al. [14] reported that smooth muscle cell apoptosis correlated with the amount of fibrillin in primary varicose veins. They compared two groups according to age but did not use any antibody for elastin or serial sections. In the present study, lumen blockage by fibrillin-1-rich tissue and weakened α-SMA expression occurred simultaneously (Figs 3 and 4). However, further studies are required to identify the mechanism underlying lumen blockage and reduction in α-SMA expression. A limitation of the study was that, without ascorbic acid, the blockage of the blood vessel lumen was observed in two of four blood vessels, but the cause of the blockage in two blood vessels was unclear. The bovine blood vessel model developed in this study is useful for investigating damaged blood vessels and lumen blockage. This in vitro model incorporates vascular endothelial cells and smooth muscle cells, as well as the intima, media, and adventitia of blood vessels, and investigation of this model will help study clinical vascular diseases.

## Abbreviations

(FBXW2): F-box and WD-40 domain-containing protein 2
(α-SMA): alpha-smooth muscle actin

## Data Availability

A pre-print version of this article is available on bioRxiv (https://doi.org/10.1101/2024.02.06.579234). The data that support the findings of this study are openly available in [Biomimetics in Fig.2 a, b 2022-12] at [DOI: 10.3390/biomimetics8010007], reference number [10].

## Acknowledgments

This work was supported by JSPS KAKENHI Grant Number 21K09947. I would like to thank Editage (www.editage.jp) for English language editing, Kobe Chuo Chikusan for providing the bovine legs, and KAC Co., Ltd. for the serial sections.

## Author Contributions

M. Akiyama designed research, performed immunohistochemistry, analyzed data, and wrote the paper. There is no co-author.

